# Identification of Novel Marine-based Inhibitors against Tetracycline Destructase in *Acinetobacter Baumannii* using Computational Approaches

**DOI:** 10.1101/2025.06.08.658539

**Authors:** Sathish Kumar Marimuthu, Pratima Chockalingam, Vaishnavi Nagarajan, Subbiah Thamotharan, Vigneshwar Ramakrishnan

## Abstract

Tetracyclines are indispensable antibiotics employed to treat a broad spectrum of bacterial infections. However, the emergence of clinical pathogens exhibiting significant resistance to these drugs has posed a formidable challenge in managing bacterial diseases effectively. This resistance is attributed to the proliferation and diversity of resistance genes and various mechanisms that render tetracyclines inactive. A prevalent mechanism of resistance is enzymatic inactivation by tetracycline destructases, which compromises the efficacy of tetracyclines. Consequently, there is a pressing need to discover novel molecules capable of inhibiting tetracycline destructases activity and thereby enhancing the drug’s potency. In this study, we identify novel inhibitors for tetracycline destructase (TDase) in *Acinetobacter Baumannii*, utilizing compounds derived from marine environments. Through a high-throughput virtual screening approach, we have investigated the potent natural marine compounds interaction and dynamic behaviour of tetracycline destructase through molecular simulations. These findings hold promise for the development of novel and efficacious therapeutics against bacterial infections.

## Introduction

*Acinetobacter Baumannii* is a nonfermenting gram-negative bacteria that poses a significant public health threat. It is most commonly associated with patients in the intensive care unit and is associated with several clinical conditions including ventilator-associated pneumonia, bacterimia, endocarditis etc [1, 2]. The emergence of multi-drug resistant and extensively drug-resistant strains only add to the treatment challenges. It is estimated that ∼15% of the deaths attributed to AMR in patients greater than 5 years old are from *Acinetobacter Baumannii* [3]. Considering these, the World Health Organisation has listed *A. baumannii* as one of the critical priority bacteria [4]. Severe Multi-Drug Resistance–*Acinetobacter Baumannii* and Carbepenem Resistance-*Acinetobacter Baumannii* infections are treated with the last-resort antibiotic Colistin, but its use is limited due to its neurotoxic and nephrotoxic effects [5]. Tigecycline shows bacteriostatic activity which is attributed to the inhibition of bacterial protein synthesis by blocking aminoacyl-tRNA attachment at the A site of the 30S ribosome [6]. A high prevalence of multiple heteroresistance to tigecycline and colistin was observed among *A. baumannii* clinical isolates [7]. Resistance to tetracyclines is, in general attributed to efflux pumps and ribosomal protection proteins. However, recent studies show the emergence of a third mechanism, the tetracycline inactivating enzymes, the Tetracycline Destructases [8, 9]. Tetracycline Destructases (TDases) are categorized into two types: type 1 Tet(X)-like TDases, which are commonly found in clinical pathogens, and type 2 soil-derived TDases. The structure of the TDases in general consists of the FAD-binding domain and the substrate binding domain connected by a C-terminal bridge helix [10]. FAD cofactor is essential in type 1 tetracycline destructases (TDases) acting as a redox cofactor to oxidize the substrates. Throughout the catalytic cycle, FAD undergoes reversible redox reactions, transitioning between states (“IN” and “OUT” orientations) to aid in substrate binding, oxidation, and regeneration [11]. Type 1 TDases are known to inactivate even the third-generation tetracyclines such as tigecycline thus making it an important clinical threat [12]. To-date more than ten Tet(X) variants have been reported across various bacteria and has thus emerged as an important target in tackling tetracycline resistance [11].

Marine metabolites have now emerged as an important source of novel compounds with therapeutic properties. For example, computational screening of marine metabolites identified potent inhibitors of dipeptidyl peptidase-4 (DPP-4), an important target in the treatment of Type 2 Diabetes Mellitus [12]. Marine compounds with potential for inhibiting acetylcholinesterase have been identified, highlighting their possible applications in treating neurodegenerative diseases such as Alzheimer’s [13]. Recent research also indicates that marine natural compounds have antimicrobial activity and thus can serve as new scaffolds of antibiotics. Vitroprocines, antibiotics, effective against *Acinetobacter Baumannii*, were identified from marine *Vibrio sp*. [14]. A high-throughput screening of the Comprehensive Marine Natural Products Database (CMNPD) also revealed potent inhibitors of 5-enolpyruvylshikimate-3-phosphate (EPSP) synthase in *Acinetobacter Baumannii* [15]. Similarly, marine compounds that may inhibit Metallo-beta-lactamases, specifically NDM-1, VIM-2, and IMP-1, were identified highlighting the potential of marine natural products as a valuable resource for developing novel antimicrobial agents [16].

In this research, we explored natural marine products as lead compounds to investigate their potential as inhibitors of tetracycline destructase (TDases) and as potent alternatives to 5a,6-anhydrotetracycline. We conducted molecular docking simulations to evaluate the binding affinity and interactions of various natural marine compounds with the active site of Tet(X) from *A. baumannii*. Promising compounds were then subjected to molecular dynamics simulations to examine the stability of the protein-ligand complexes over extended time periods. Our results indicate that several natural compounds exhibit favourable binding interactions with the active site of Tet(X), suggesting their potential as inhibitors. Additionally, the simulations revealed the dynamic behavior of the active site residues. This computational study provides valuable insights into the potential of natural marine compounds as TDase inhibitors. The identified compounds could serve as promising candidates for further experimental validation and optimization as novel therapeutics in the fight against antibiotic resistance.

## Materials and Methods

### Protein Preparation

The sequence of tetracycline inactivating monooxygenase (Tet(X)) in *A. baumannii* was retrieved from NCBI with accession number: WP_236754017.1. The three-dimensional structure of this sequence was obtained by homology modeling using SWISS-MODEL [17]. The X-ray crystal structure of Tet(X6) from *Escherichia coli* (PDB ID: 8ER0) which has a sequence identity of 99.74% and coverage of 100% served as the template for building the model. The modeled structure was validated using Ramachandran plot. The protein preparation wizard in Maestro [18], was then used to pre-process the protein structure followed by optimizing hydrogen bonds, and energy minimization (the RMSD cutoff of heavy atoms is 0.30 Å) using the OPLS4 force field [19].

### High throughput Virtual Screening

The Comprehensive Marine Natural Product Database (CMNPD), containing 47,451 compounds, was chosen as the ligand database [20]. The ligands were prepared using the LigPrep module (Schrödinger, LLC, New York). These prepared compounds were then docked using the Virtual Screening Workflow (VSW), with the top hits selected from Glide’s Extra Precision (XP) mode [21]. Finally, the ligand interaction profiles of the top three marine natural compounds were analyzed to identify the most promising candidates.

### Molecular dynamics simulations of high-scoring compounds

Molecular dynamics (MD) simulations of the three protein-ligand complexes with the highest binding scores were done using the Desmond module available in the Schrodinger software. In addition, MD simulation of the protein-bound with anhydrotetracycline was also done [22]. Each complex was placed inside a cubic box whose wall is at a distance of 10 Å from the outermost protein atom. Each system was solvated and ions were added randomly within the box to neutralize the system. TIP3P model was used to represent water and the OPLS4 forcefield was used to represent the protein, ligands, and ions. The systems were subjected to multiple phases of NPT ensemble relaxation process. The final production MD was carried out for 100 ns, maintaining constant pressure (1.01325 bar) and temperature (300 K). RMSD, RMSF were plotted using XMgrace [23]. Additionally, post-MD MM-GBSA (Molecular Mechanics with Generalised Born and Surface Area solvation) free energy analysis was performed for each of the four protein-ligand complexes for the last 50 ns simulations.

### Principal component and free energy landscape (FEL) analyses

Principal Component Analysis (PCA) is a widely utilized method for examining the collective movements of residues in proteins [24, 25]. Principal Component Analysis (PCA) was performed using data from the last 60 ns of the MD simulation. This approach involves constructing a covariance matrix based on the positional deviations of the C_α_ atoms from the trajectory. The eigen decomposition of this matrix yields eigenvectors arranged in decreasing values of corresponding eigen values. While the eigen vectors denote the directions of motions, the corresponding eigenvalues measure the extent of the motions (variations). The first eigenvector, associated with the highest variance, is termed Principal Component 1 (PC1), followed by subsequent eigenvectors with decreasing variances, named Principal Component 2 (PC2), Principal Component 3 (PC3), and so on. To quantify the conformational distributions of the simulated systems, we used the Gibbs Free Energy Landscape (FEL) calculated against the first two principal components (PC1 and PC2). The Gibbs free energy is derived from the probability distribution of the projections of the trajectory structures along these principal components using the following equation:

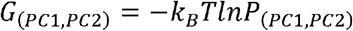

Where *k*_*B*_ is the Boltzmann constant,

T is the temperature and

*P*_*(PC1,PC2)*_ is the normalized joint probability distribution.

### Dynamic Cross-Correlation Matrix

The dynamic cross-correlation matrix, which contains the correlation coefficients for all pairs of residues, is represented by the following equation:

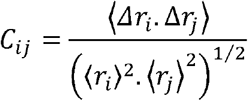

where *i* and *j* represent two *C*_*α*_ atoms of different residues, and Δr_*i*_ and Δr_*j*_ are their displacement vectors. Angle brackets denote the ensemble average [26, 27, 28]. A positive C(*i,j*) value indicates positively correlated movements, where both atoms move in the same direction, while a negative value signifies anti-correlated movements, where the atoms move in opposite directions. C(*i,j*) values of 1 and −1 denote completely correlated (blue) and anti-correlated (red) motions, respectively. White color is used for C(*i,j*) values between −0.5 and 0.5 to create a clear 2D plot. Off-diagonal colors indicate residue motions with strong correlations (0.51 to 1.0) and anti-correlations (−0.51 to −1.0). A dynamic cross-correlation matrix was performed using data from the last 60 ns of the MD simulation.

## Results and Discussion

### High-throughput virtual screening (HTVS) of marine compounds

The three-dimensional structure of *Acinetobacter Baumannii* tetracycline-inactivating monooxygenase Tet(X) was modeled and validated using Ramachandran plot. The modeled structure with flavin-adenine dinucleotide (FAD) (in white and blue) and 5a,6-anhydrotetracycline (TDC) (in orange) (Fig.1). Fig.1 box highlights the residues around 4 Å of 5a,6-anhydrotetracycline which were used as binding site for the virtual screening of marine compounds. Marine natural compounds were downloaded from the Comprehensive Marine Natural Product Database (CMNPD). High-throughput virtual screening (HTVS) was performed and the compounds with top three binding score were identified through extra precision docking. The binding score and interacting residues are tabulated in Table 1. From the results we find that all the three compounds have a binding score higher (in the negative scale) than that of anhydrotetracycline. However, the MM-GBSA binding energy is indeed highest (in the negative scale) for anhydrotetracycline.

**Table 1.**
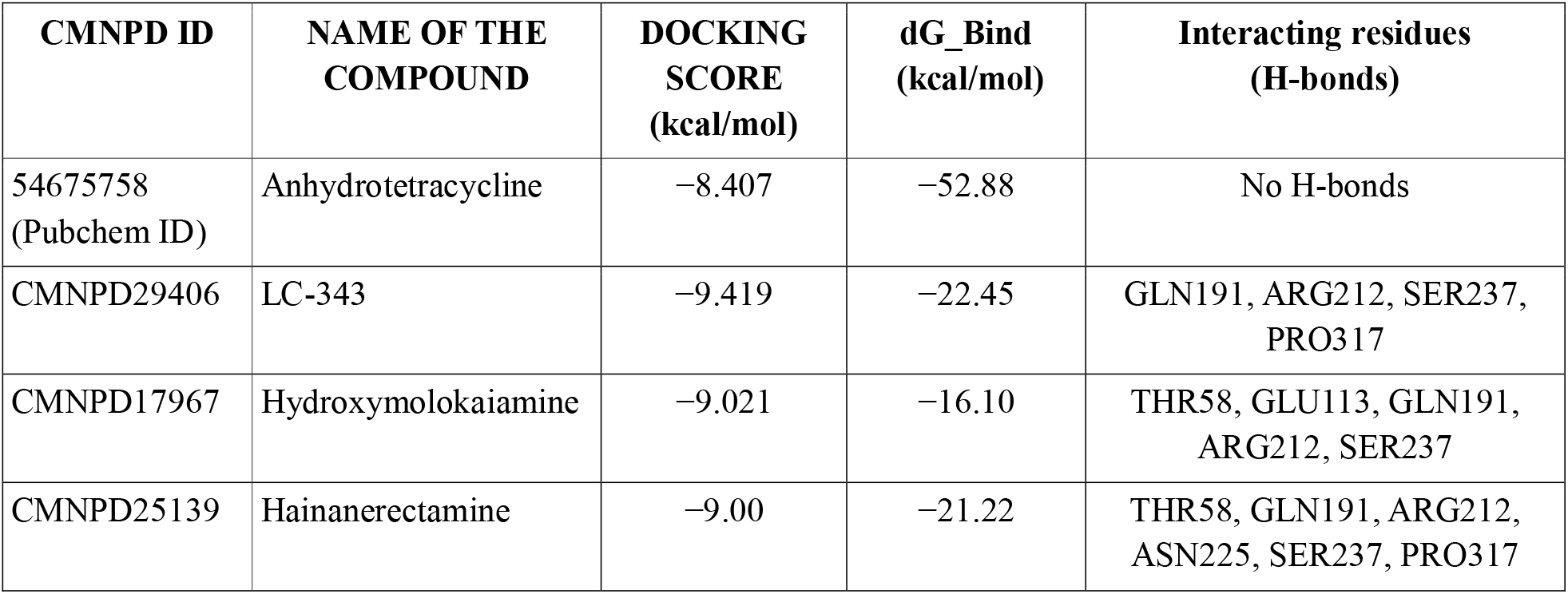
Docking scores of the top hits along with their interacting residues.

**Fig. 1.**
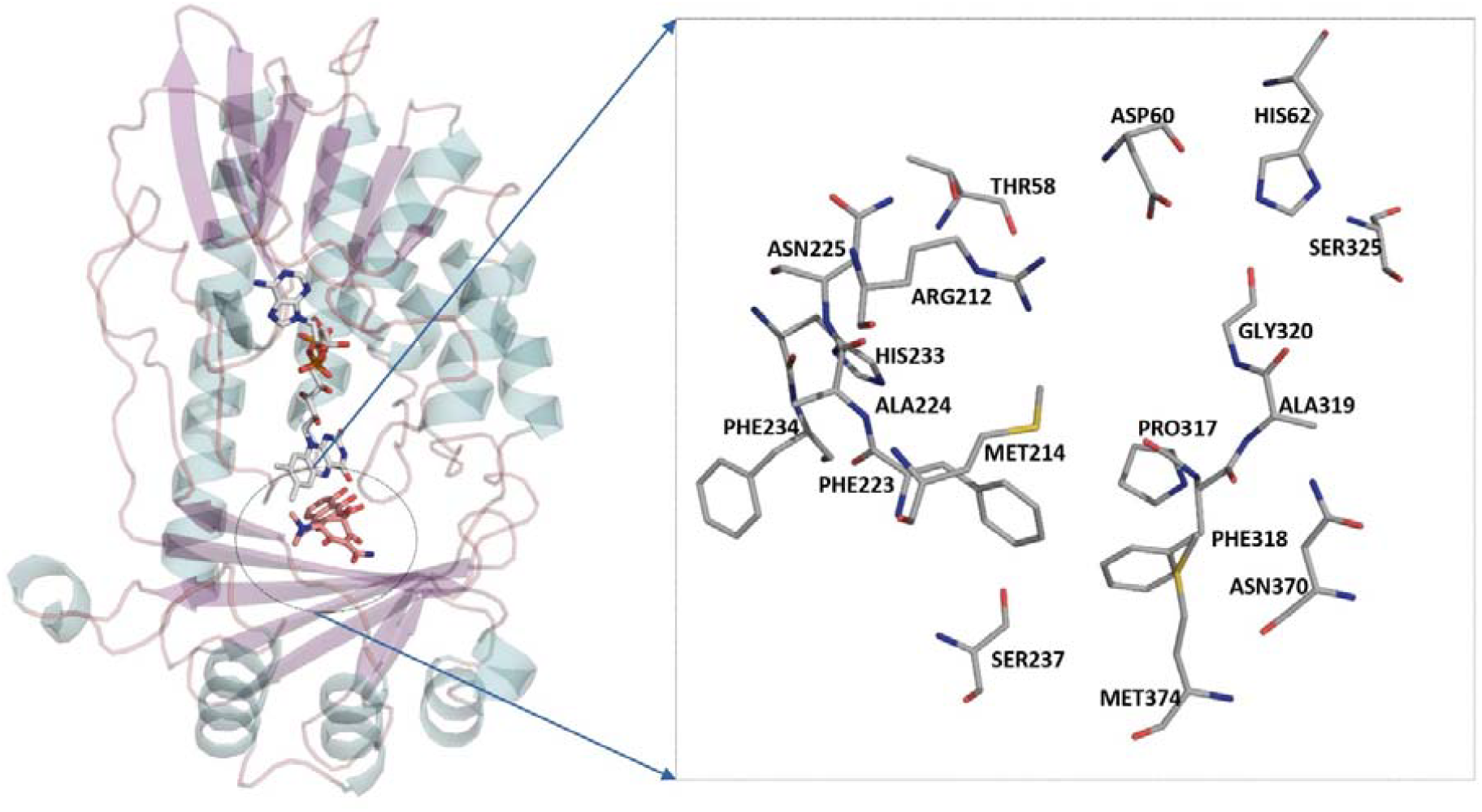
Three-dimensional structure of modeled tetracycline-inactivating monooxygenase Tet(X) from *Acinetobacter Baumannii* bound to with flavin-adenine dinucleotide (white and blue) and 5a,6-anhydrotetracycline(orange), and the dashed box indicates the residues within 4 Å of anhydrotetracycline.

### Drug-like properties of the top-hit compounds

The top hit compounds identified from the CMNPD database (CMNPD29406, CMNPD17967, CMNPD25139), which target the tetracycline-inactivating monooxygenase Tet(X) in *Acinetobacter Baumannii*, were further evaluated for drug-like properties using the QikProp module of the Schrodinger Suite (2023-2). The QikProp drug-like properties of these compounds are detailed in Table 2. The results indicate that the drug-like properties of the top ligands fall within the acceptable range for most QikProp parameters. Additionally, toxicity predictions for these compounds were assessed using the PROTOX web server [29]. Table 3 reveals that two of the lead molecules fall into toxicity class 3, while one is classified as class 5. Both classes are considered toxic, with class 5 potentially harmful if ingested. Despite this, the identified molecules hold promise as lead compounds for further optimization.

**Table 2.** Qikprop properties of top hits compounds.

| Compound name | QPlogPw | QPlogS | QPPCaco | QplogBB | % HOA | Rule of five | Rule of three |
| --- | --- | --- | --- | --- | --- | --- | --- |
| CMNPD29406 | 16.201 | – 1.632 | 24.020 | – 1.973 | 53.719 | 0 | 1 |
| CMNPD17967 | 12.297 | – 0.166 | 15.964 | – 0.413 | 51.207 | 0 | 1 |
| CMNPD25139 | 7.916 | – 2.659 | 125.353 | – 1.594 | 72.270 | 0 | 0 |

**Table 3.** Toxicity prediction of top hits compounds.

| Compound name | LD50 (mg/kg) | Predicted Toxicity Class | logP | Molweight | Prediction Accuracy |
| --- | --- | --- | --- | --- | --- |
| CMNPD29406 | 300 | 3 | – 2.83 | 302.32 | 67.38 % |
| CMNPD17967 | 2300 | 5 | 3.33 | 368.07 | 68.07 % |
| CMNPD25139 | 200 | 3 | 1.4 | 247.25 | 67.38 % |

**Toxicity prediction class** description:

**Class 1:** LD50 ≤ 5 – Fatal if swallowed.

**Class 2:** 5 < LD50 ≤ 50 – Also fatal if swallowed.

**Class 3:** 50 < LD50 ≤ 300 – Toxic if swallowed.

**Class 4:** 300 < LD50 ≤ 2000 – Harmful if swallowed.

**Class 5:** 2000 < LD50 ≤ 5000 – May be harmful if swallowed.

**Class 6:** LD50 > 5000 – Non-toxic.

### Stability of the complexes

The RMSD of the protein *C*_*α*_ atoms for the four systems relative to their initial structures over the 100 ns simulation period (Fig. 3A). From the RMSD plot, we find that CMNPD29046 and CMNPD25139 complexes were more stable compared to other systems. Specifically, the CMNPD29046 and CMNPD25139 complex systems showed stable equilibration from 40 ns to 100 ns. A slight deviation was observed in the TDC and CMNPD17967 complexes starting from 75 ns. The average RMSD value was calculated for all four systems throughout their entire simulations. The average RMSD value for the TDC system was found to be 2.48 Å. The corresponding average RMSD values for the other systems were as follows: 2.15 Å for CMNPD29406, 2.56 Å for CMNPD17967, and 2.25 Å for CMNPD25139.

**Fig. 2.**
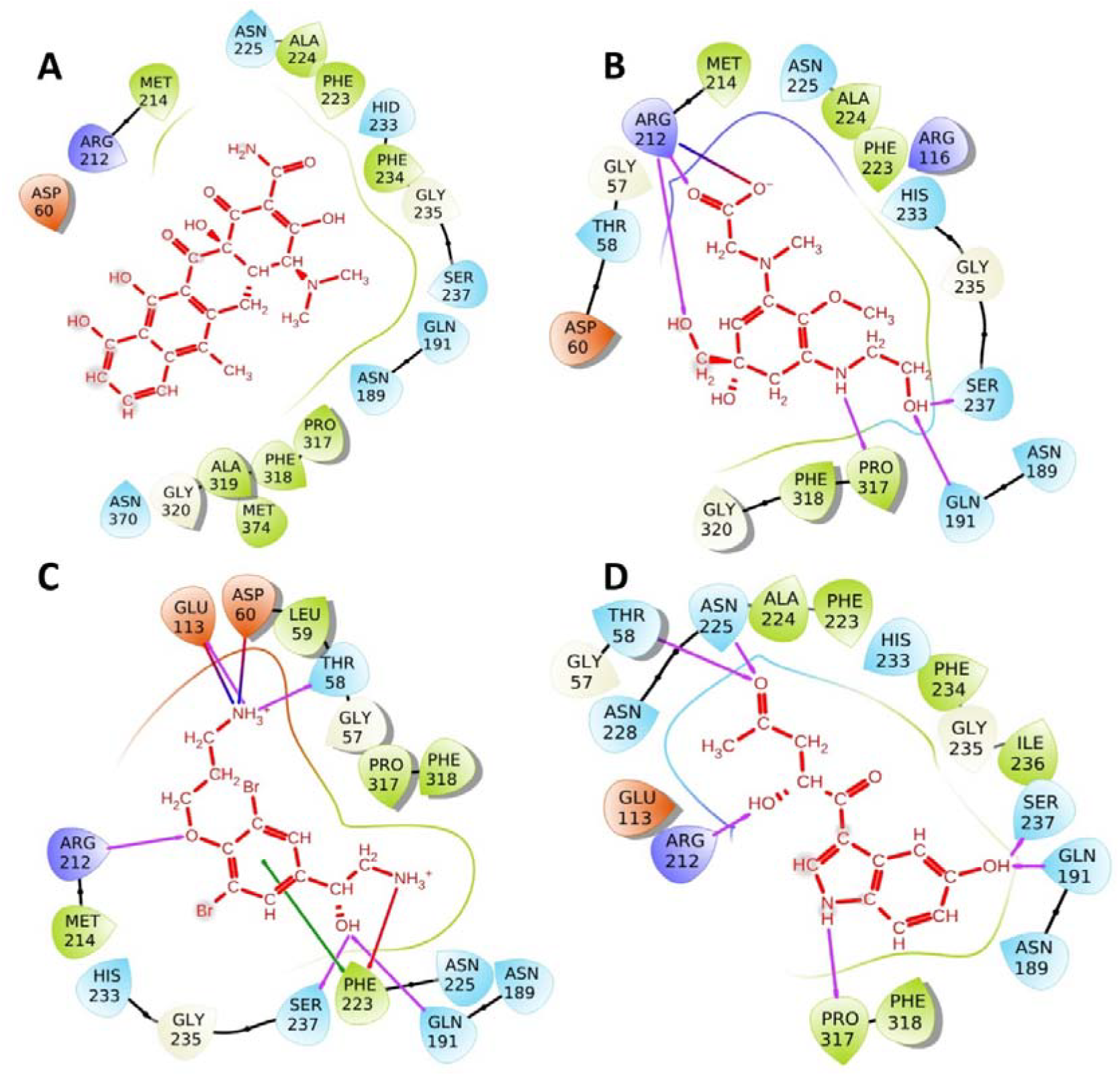
Ligand interactions of the top hits from HTVS against tetracycline-inactivating monooxygenase (A). Anhydrotetracycline, (B). CMNPD29406, (C). CMNPD17967, (D). CMNPD25139

**Fig. 3.**
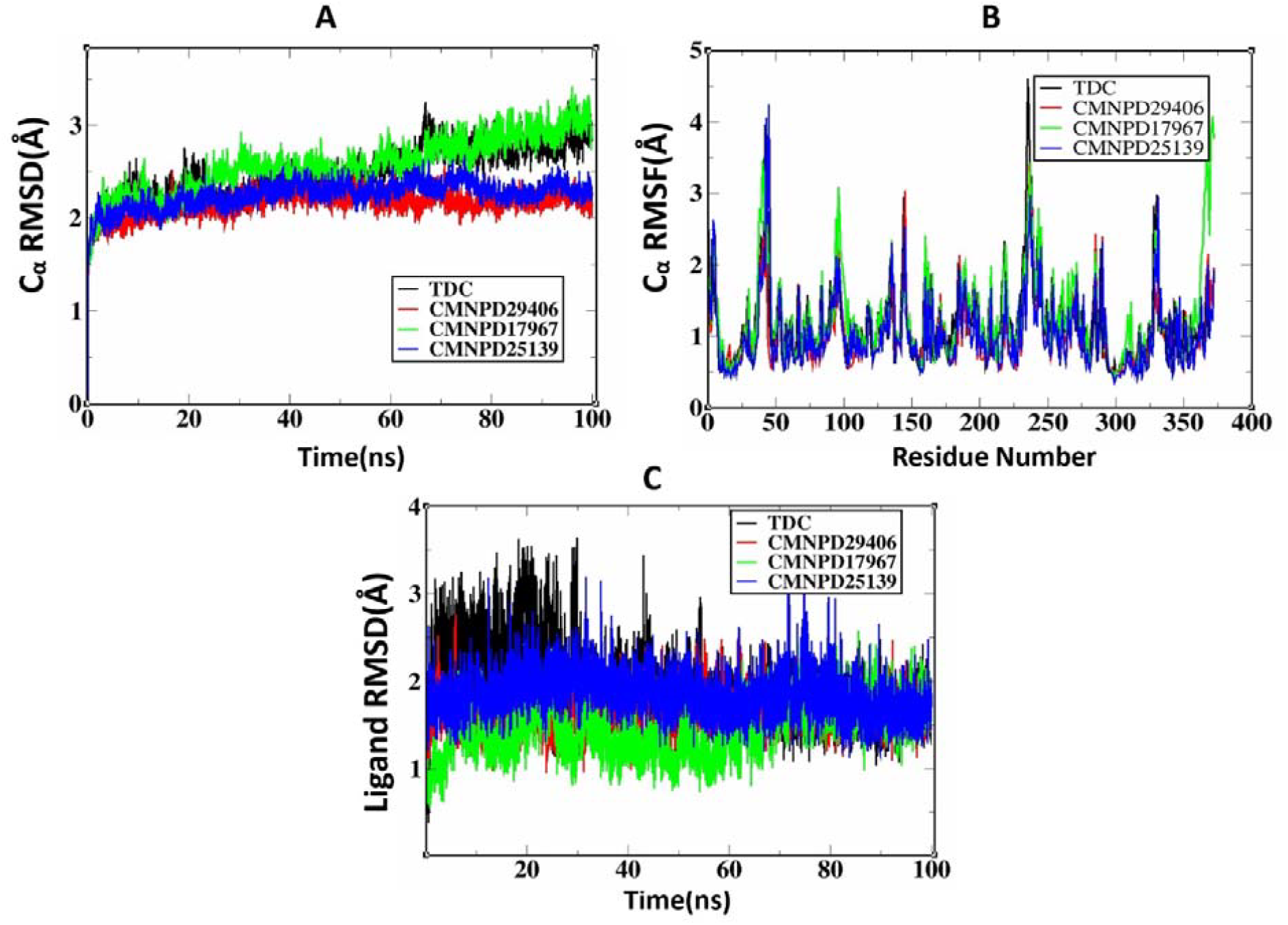
(A) C_α_ RMSD of the four protein-ligands complexes (Anhydrotetracycline, CMNPD29406, CMNPD17967, CMNPD25139). (B) C_α_ RMSF of the four protein-ligands complexes (Anhydrotetracycline, CMNPD29406, CMNPD17967, CMNPD25139). (C) RMSD of the four ligands (Anhydrotetracycline, CMNPD29406, CMNPD17967, CMNPD25139).

The RMSF of the four complexes, showing similar residue fluctuation behavior (Fig. 3B). Residues involved in binding (THR58, ASP60, HIS62, GLN191, ARG212, PHE223, ALA224, ASN225, HIS233, PHE234, MET214, GLY325, SER237, PRO317, PHE318, ALA319, GLY320, ASN370, MET374) do not exhibit higher fluctuations in the RMSF plot. Conversely, residues such as ARG53, PHE55, SER246, LYS247, LYS300, GLU106, ASN107, ARG249, GLN253, VAL43, TYR44 which are in the loop regions distant from the active sites how higher fluctuation. The Root Mean Square Deviation (RMSD) values for the ligands across different systems (Fig. 3C). We also calculated the binding free energy for each of the complexes using the snapshots from the MD simulations. The results show that post MD-MM/GBSA analysis indicates that the ΔG_bind value for TDC is −61.36 ± 0.37, while the remaining three marine natural compounds have ΔG_bind values of −49.63 ± 0.55 (CMNPD29406), −49.87 ± 0.39 (CMNPD17967), and −53.60 ± 0.37 (CMNPD25139). Taken together, these results indicate that the systems remain stable over the 100 ns simulation trajectory.

### Ligand-binding mode

We conducted a thorough analysis of protein-ligand interactions that persisted for more than 30% of the simulation time to identify crucial binding residues. The residues that maintained contact with the ligand in over 30% of the frames (Fig. 4). These interactions are significant and play a key role in stabilizing the ligand at the active site. In general, the binding residues THR58, ASP60, GLN191, ARG212, PHE223, ALA224, ASN225, HIS233, PHE234, GLY325, SER237, PRO317, PHE318, ALA319 are involved in establishing direct interactions and as well water-mediated interactions throughout the simulations. In the anhydrotetracycline complex (Fig. 4A), the ligand exhibits direct interactions with ARG212, HIS233, PHE223, and ASN225. Additionally, ALA224 and THR58 show water-mediated interactions with the protein during the simulation. In the CMNPD29406 system (Fig. 4B), ARG116 and ALA319 participate in water-mediated interactions. Additionally, residues ARG212, ASN225, and THR58 establish direct contacts with CMNPD29406. In CMNPD17967 complex system (Fig. 4C), only ILE236 and ILE190 have water mediated interaction and other residues (ASP60, GLU113, THR58, ASN225, PRO317, PHE318, GLN191, PHE223) have direct contacts with the ligand. The residues ASN189, PHE223, GLU113 have direct contacts and remaining residues (GLY57, HIS233, PRO317) having water mediated interactions. In CMNPD25139 complex system (Fig. 4D), three water mediate interactions found with the residues GLY57, GLN191 and PRO317 and direct interactions with HIS233, GLN191, ASN189 and GLU113.

**Fig. 4.**
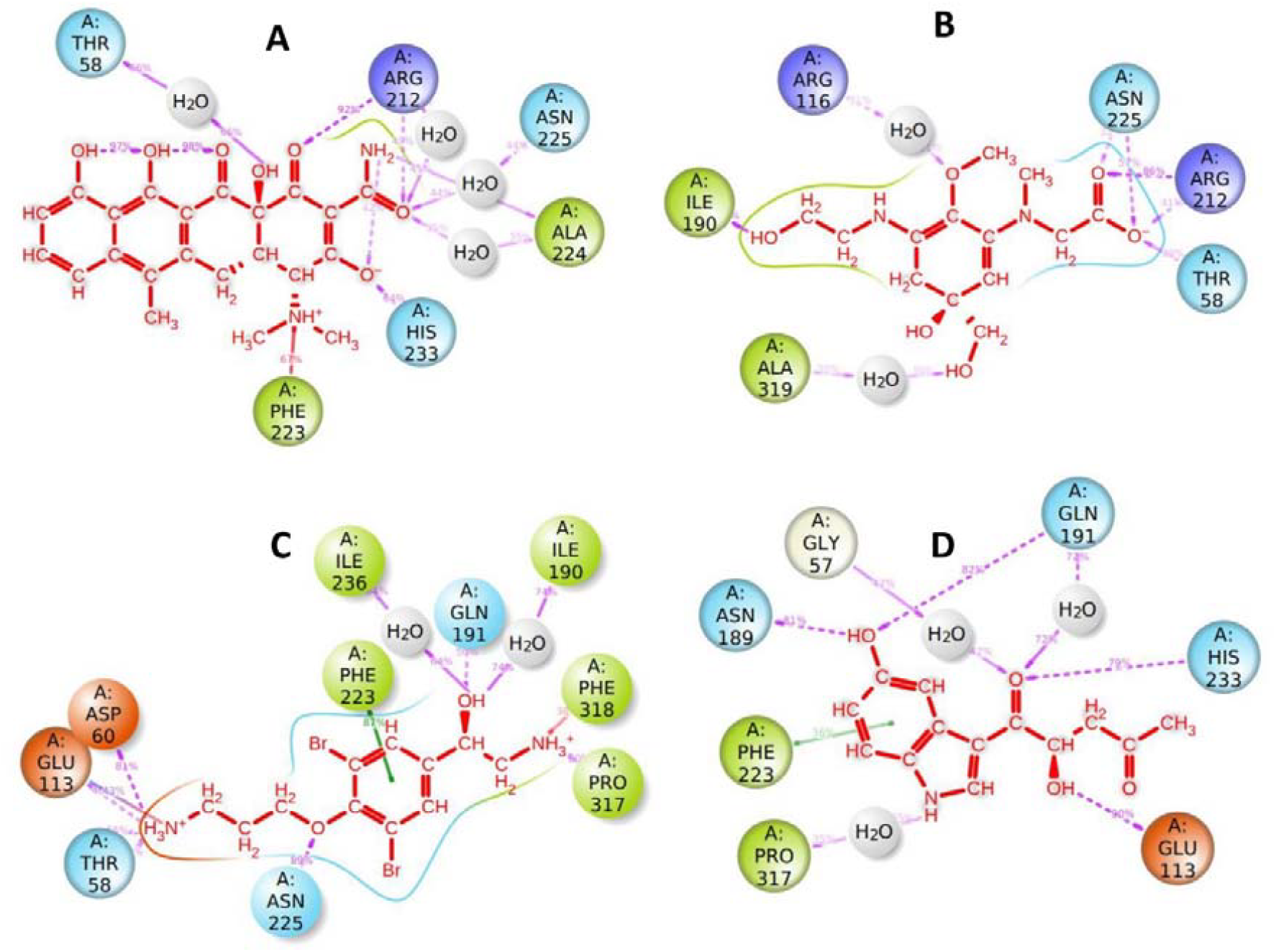
Ligand protein contacts during (A). Anhydrotetracycline, (B). CMNPD29406, (C). CMNPD17967, (D). CMNPD25139

### Dynamics Cross-Correlation matrices analysis

To further understand how each of the ligands affect the protein dynamics, we calculated the dynamic cross-correlation matrices (based on C_α_ atoms) which reveal correlated and anti-correlated motions. The dynamic cross-correlation matrix plot for the four simulated systems (Fig. 5). In this plot, correlations are depicted in blue color, and anti-correlations appear in red. In general, we observed that CMNPD29406 and CMNPD25139 systems exhibit fewer anti-correlated motions among their residues compared to TDC and CMNPD17967 systems. In the TDC complex system, specific residues within the FAD binding sites display patterns of anticorrelation with other residues. Specifically, residues 148, 149, and 150 within the FAD binding sites show anticorrelation with residues 249 to 259, which are highlighted as C. Similarly, residues 178, 179, and 180 within the FAD binding sites exhibit anticorrelation with residues 329 to 338, marked as D. However, in contrast to the TDC complex system, these anticorrelation patterns are not observed in the CMNPD29406 and CMNPD25139 systems. In the TDC system, residues 250 to 285 (highlighted in the box A) exhibit anti-correlated movements with residues 107 to 141. These particular residues are not involved in the active site of the protein. Similarly, these regions show less anti-correlation in the CMNPD29406 and CMNPD25139 systems. Residues 117 to 137 exhibit an anticorrelation in the TDC and CMNPD17967 systems. However, this negative correlation is not observed in the CMNPD29406 and CMNPD25139 systems and the residues found in both A and B are not part of the protein’s binding sites.

**Fig. 5.**
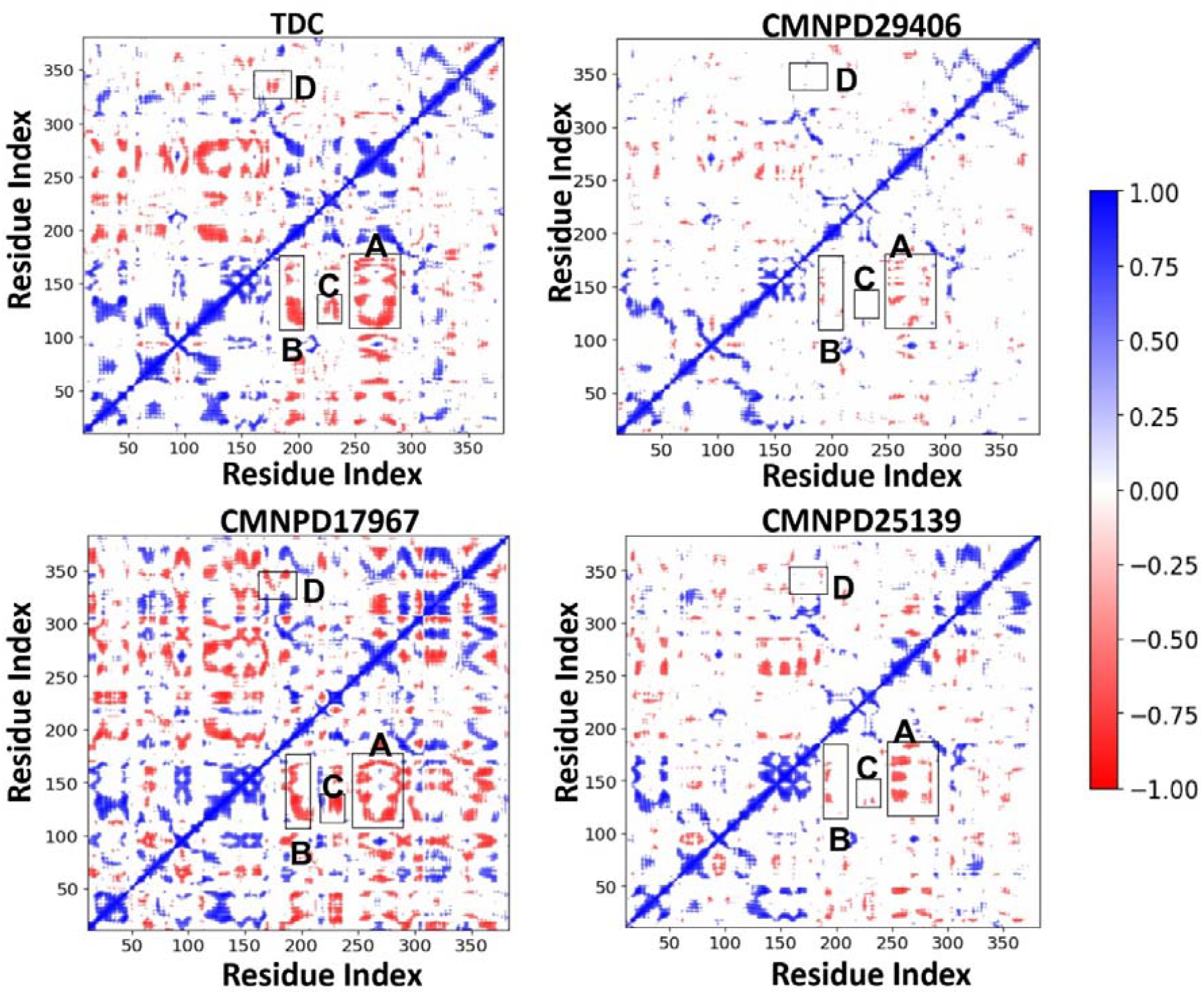
Dynamic cross-correlation matrix plot depicts correlations in blue and anti-correlations in red for Anhydrotetracycline (TDC), CMNPD29406, CMNPD17967, and CMNPD25139.

### Principal Component Analysis (PCA) and Free Energy Landscape (FEL) analysis

Principal component analysis was performed to investigate the dominant motions and conformational states of the protein in the four systems. The principal component analysis reveals that the first ten principal components account for a significant portion of the overall motion in the four complex systems, with contributions of 57.52% for the Anhydrotetracycline (TDC) system, 49.54% for the CMNPD29406 system, 59.67% for the CMNPD17967 system, and 49.53% for the CMNPD25139 system. The aggregated contribution of the top ten principal components is shown in Supplementary Fig. 5. To further elucidate these conformational states, we constructed the free energy landscape for each system using the first two principal components (PC1 and PC2). This approach provides a detailed view of the energy minima and the conformational diversity sampled during the molecular dynamics simulations. In all protein-ligand complexes, approximately 80% of the protein structures have an energy value between 0.00 and 5.00 kcal/mol (refer Supplementary Table 1). For further analysis of structural dynamics, we selected 10 structures from the free energy landscape for each system, specifically those with energy levels between 0 and 2.5 kcal/mol, and examined the dynamics of the binding site residues.

The 2d free energy landscape of PC1 and PC2 components for all protein-ligand complexes (Fig. 6). The ten representative structures from each system were selected based on Gibbs free energy and different coordinates of PC1 and PC2. Supplementary Fig. 1A illustrates the superimposition of the representative structures of the anhydrotetracycline-protein system (10 structures in total). The representative structures were superimposed on the local minima structure 1, with RMSD values ranging from 0.67 to 0.94 Å. Supplementary Fig. 1B shows the individual representative structures. Both the individual and superimposed structures reveal conformational changes in the loop and alpha helix regions (residues 240-243 and 170-175), which are not near the active sites. We also analysed the interactions within the CMNPD29406-protein system and found that the hydrogen bond between ARG212 and the ligand was consistently maintained across all conformers. Additionally, the binding residues GLN191, ALA224, HIS233, and GLY320 formed hydrogen bonds in some of the conformers. Supplementary Fig. 2A illustrates the superimposition of the representative structures of the CMNPD29406-protein system (10 structures in total), with RMSD values ranging from 0.57 to 0.90 Å. Supplementary Fig. 2B depicts the individual representative structures of the CMNPD29406-protein system. Similar to previous observations, residues 240-243 and 170-175 exhibit conformational changes in the CMNPD29406-protein system. In this system, we also analysed the hydrogen bond interactions. We found that certain residues consistently maintained hydrogen bond interactions with CMNPD29406 across multiple conformers: ASN225 formed hydrogen bonds in 9 conformers, ARG212 in 8 conformers, and ILE190 in 7 conformers.

**Fig. 6.**
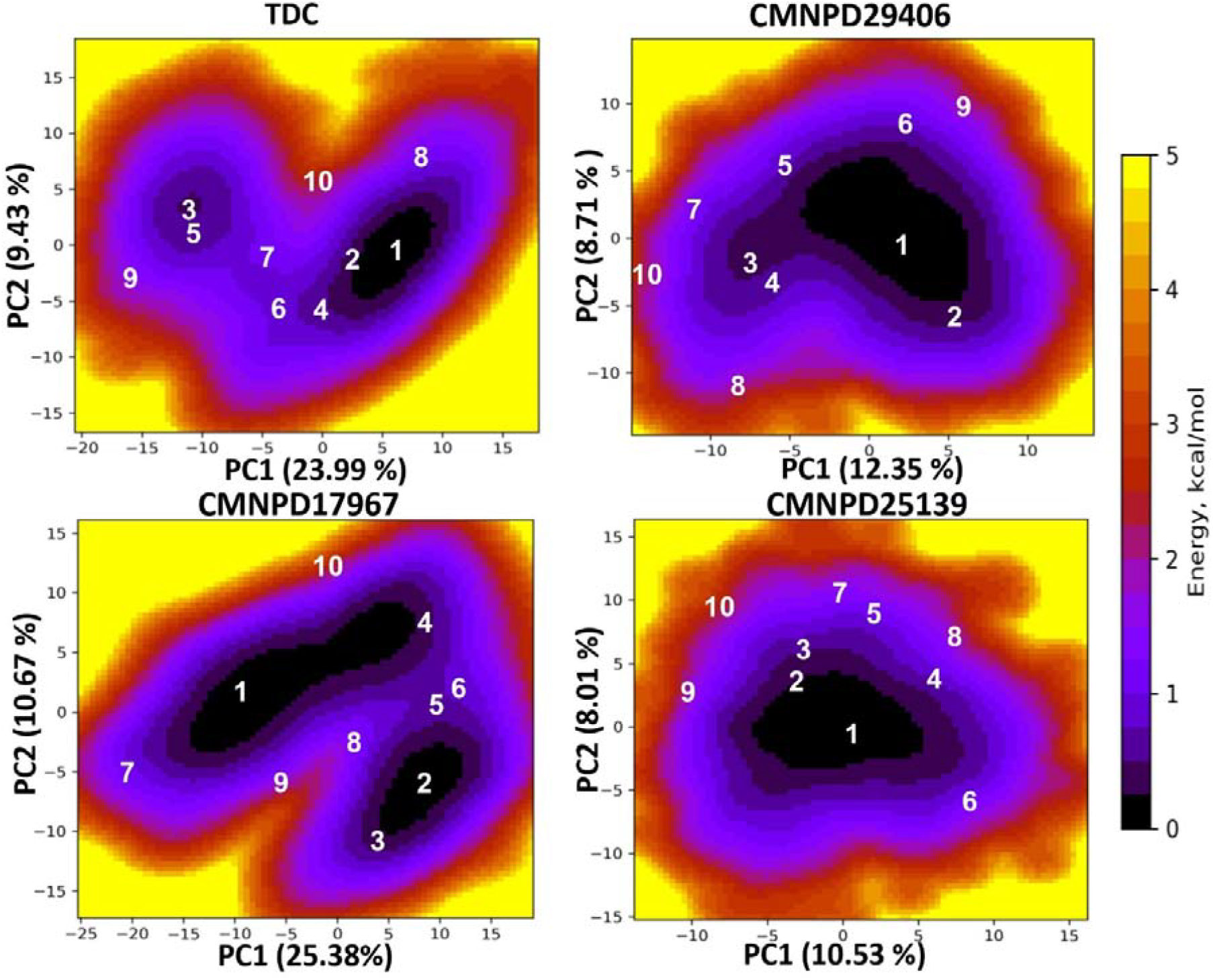
Two-dimensional free energy landscape of PC1 and PC2 components for all protein-ligand complexes.

Supplementary Fig. 3A. shows the superimposition of representative structures of CMNPD17967-protein system with the RMSD values range from 0.66 to 1.16 Å. The Individual representation of ten structures taken from PC1 and PC2 from CMNPD17967-protein system is given in Supplementary Fig. 3B. The segments from residues 240 to 243 and 170 to 175 exhibit structural conformation changes in the CMNPD17967-protein system. However, these regions are not functionally important for our study. Additionally, in the CMNPD17967-protein system, we also examined the hydrogen bond interactions. Our analysis revealed that several binding site residues consistently maintained hydrogen bond interactions with CMNPD17967 across multiple conformers. Specifically, THR58 formed hydrogen bonds in all conformers, GLN191 in 8 conformers, ASP60 and PRO317 in 10 conformers each, and ASN225 in 9 conformers. In the case of CMNPD25139-protein system, one lowest minima observed. The superimposed representative structures with the structure 1 demonstrated in Supplementary Fig. 4A. the RMSD values range from 0.69 to 0.84 Å. The Individual representation of ten structures taken from PC1 and PC2 from CMNPD17967-protein system (Supplementary Fig. 4B). The residues 170-175 shows alpha helical in 7 representative structures, remaining structures shows loop like structure and the residues 240-243 shows loop like structure in three representative structure remaining structures shows alpha helices. Based on the detailed 2D free energy landscape analysis, it was observed that residues near the binding sites (THR58, ASP60, GLN191, ARG212, PHE223, ALA224, ASN225, HIS233, PHE234, GLY325, SER237, PRO317, PHE318, ALA319) experienced minor structural changes over the 100 ns period. However, these changes did not impact ligand binding. This stability indicates that these residues play a crucial role in maintaining the ligand’s stability within the protein’s binding sites. Porcupine plots were constructed to visualize the movements of the eigenvector for each protein-ligand complexes, with the first eigenvector depicted (Fig. 7A and the second eigenvector (Fig. 7B). The loop regions surrounding the FAD and substrate, including residues 317 to 324, 44 to 47, 166 to 169, and 21 to 27, demonstrate contrasting movements in the first and second eigenvectors. The beta sheet region spanning positions 211 to 225, which is near the substrate binding sites, shows opposite motions. This dynamic behavior of the protein is largely attributed to the presence of ligand molecules.

**Fig. 7.**
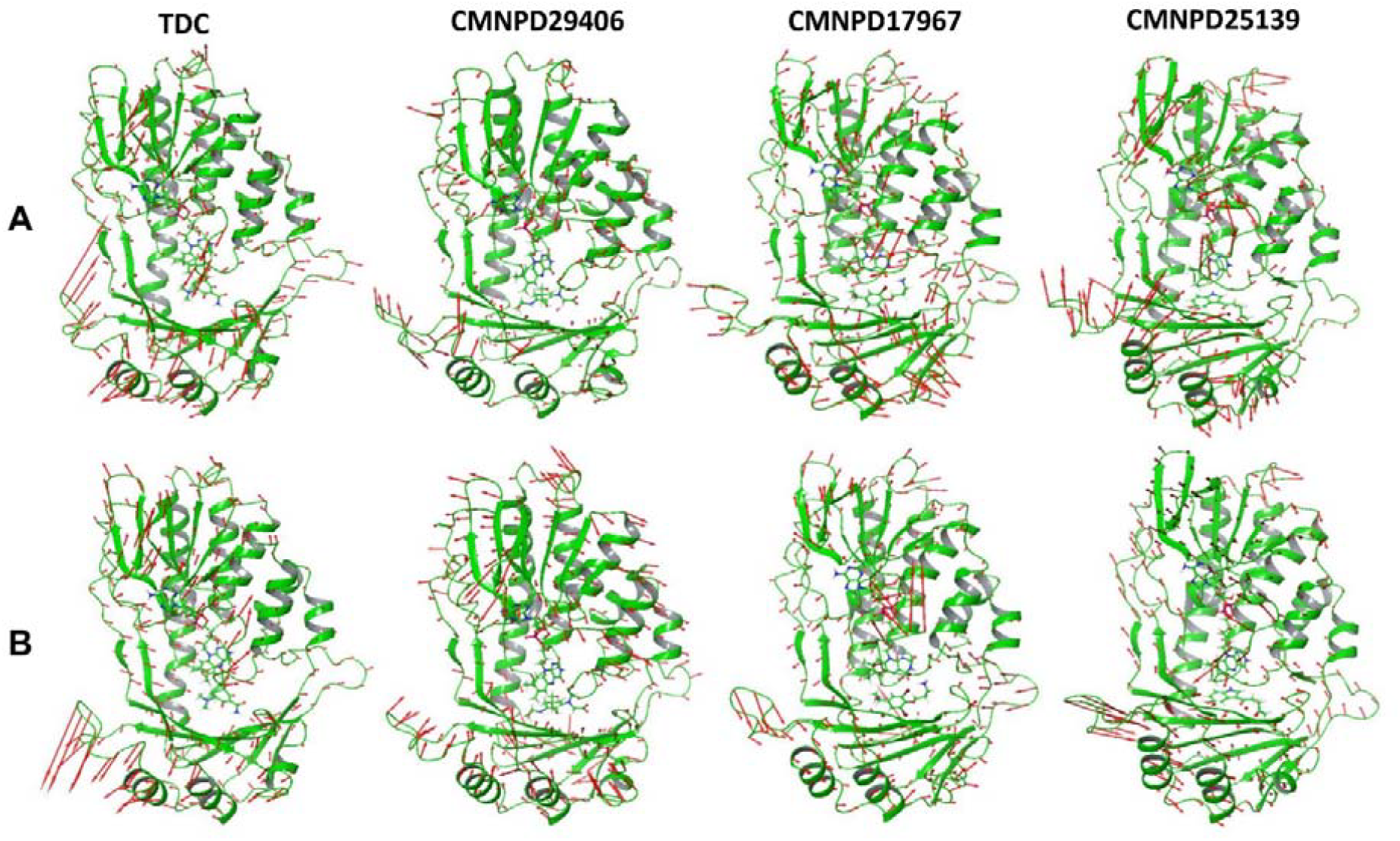
Porcupine plots to illustrate the movements of the eigenvector obtained from PCA analysis (A) first eigenvector, (B) second eigenvector

## Conclusions

Marine natural compounds have emerged as promising alternatives to traditional antibiotics, offering diverse bioactive properties that can combat drug-resistant bacterial strains. In this study, we explored marine natural products as potential inhibitors of TDases and alternatives to 5a,6-anhydrotetracycline. Using molecular docking and dynamics simulations, we identified several marine compounds with strong binding affinities to TDase active sites, indicating their potential as effective inhibitors. Specifically, CMNPD29406, CMNPD17967, and CMNPD25139 demonstrated significant interactions with key residues in TDase, suggesting potential for further therapeutic development. Our computational analyses, including molecular dynamics simulations and free energy calculations, provided insights into the stability and dynamics of these protein-ligand complexes over extended time scales. Notably, CMNPD29406, and CMNPD25139 complexes exhibited stable configurations with minimal fluctuation compared to other systems, highlighting their potential as robust therapeutic candidates. Moreover, our study identified specific binding residues involved in direct and water-mediated interactions with these marine compounds, underscoring their ability to stabilize ligands within TDase active site. These findings contribute valuable insights into the development of marine natural compounds as potential TDase inhibitors and further experimental validation and optimization in combating antibiotic resistance.

