## Supplementary File for "Identification of Novel Marine-based Inhibitors against Tetracycline Destructase in *Acinetobacter Baumannii* using Computational Approaches"

**ESI**

**Supplementary Fig 1A.** Superimposition of representative structures of TDC - protein system**. B.** Individual representation of ten structures taken from PC1 and PC2

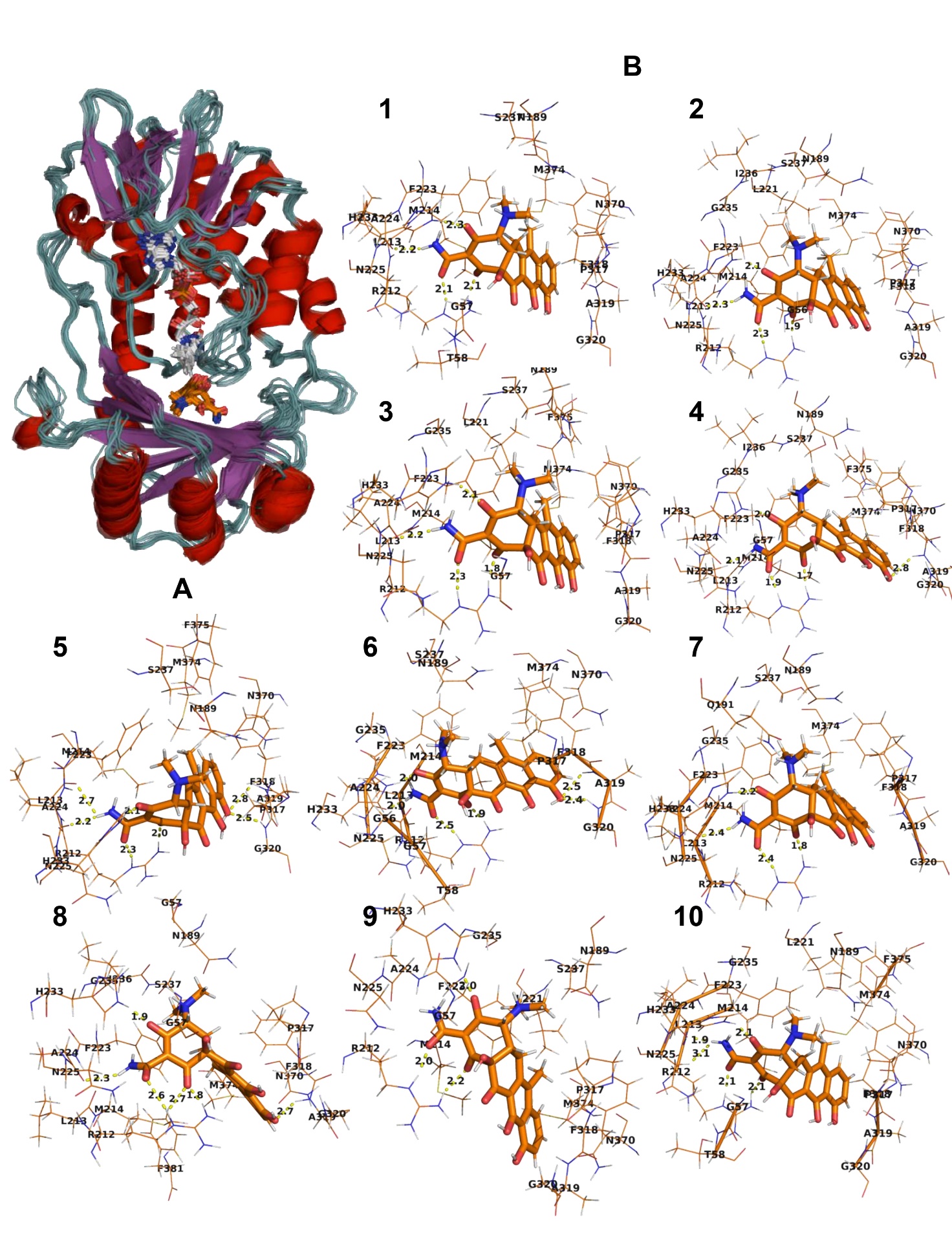

**Supplementary Fig 2A.** Superimposition of representative structures of CMNPD29406 - protein system. **B**. Individual representation of ten structures taken from PC1 and PC2

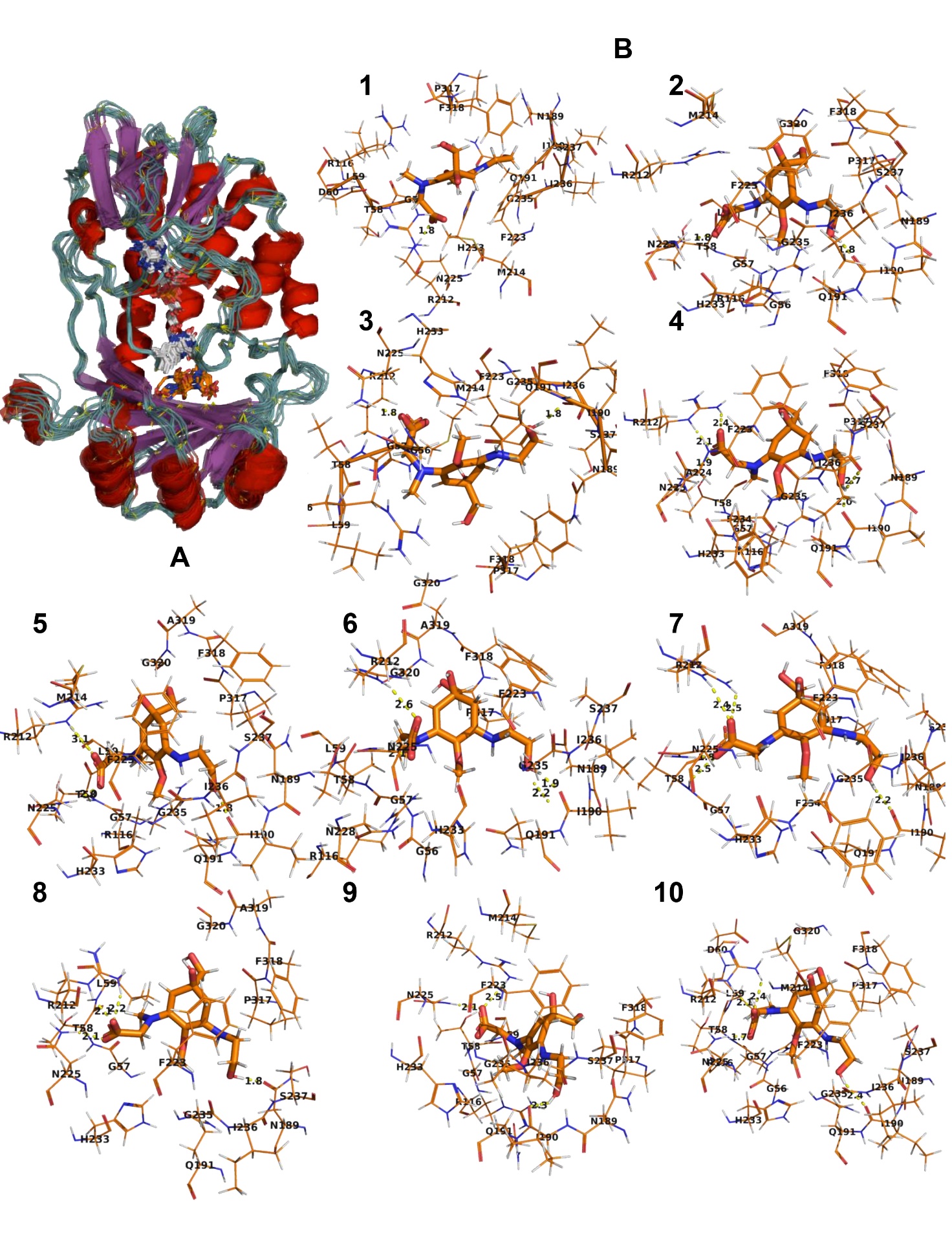

**Supplementary Fig 3A.** Superimposition of representative structures of CMNPD17967 - protein system. **B.** Individual representation of ten structures taken from PC1 and PC2

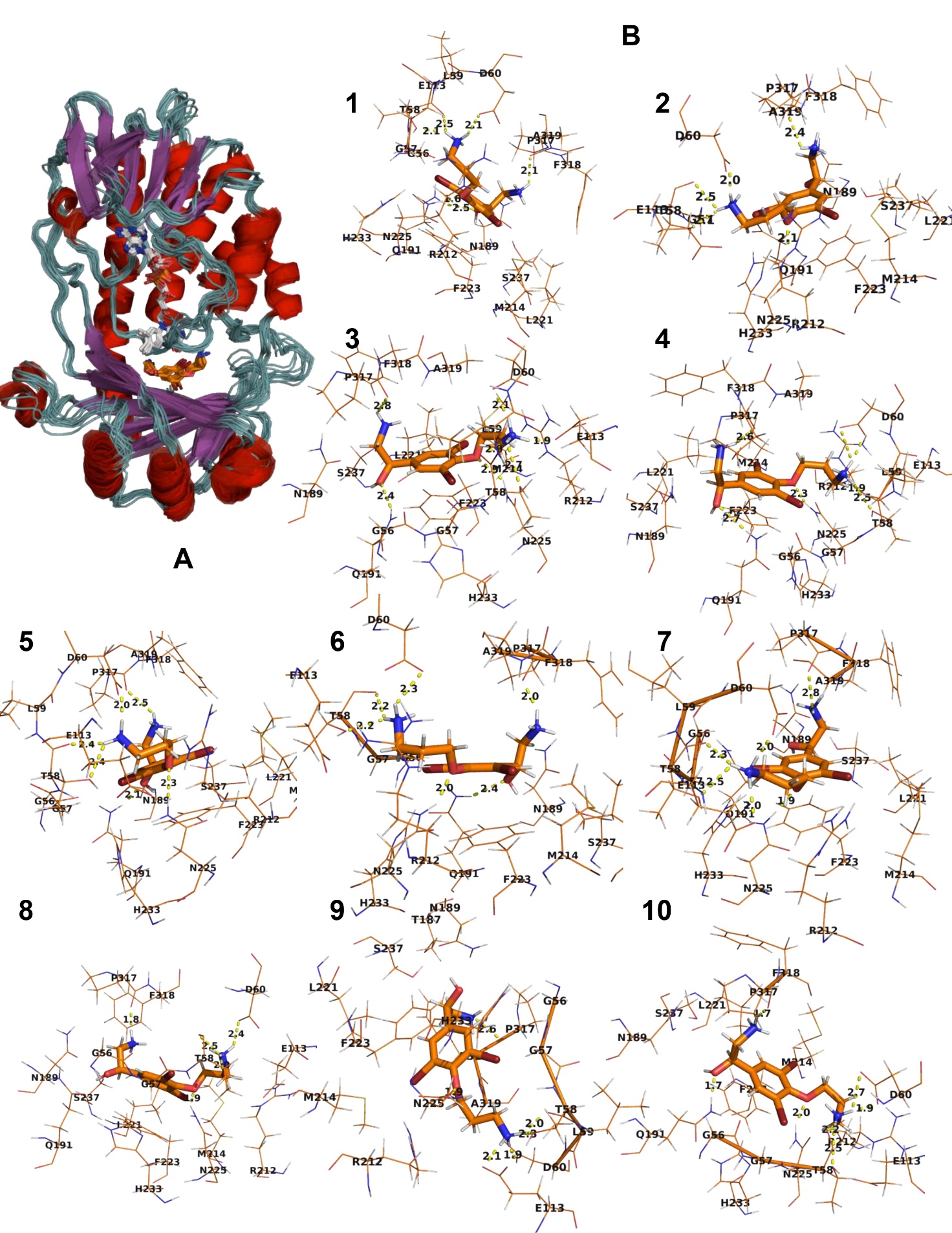

**Supplementary Fig 4.A.** Superimposition of representative structures of CMNPD25139 - protein system**. B.** Individual representation of ten structures taken from PC1 and PC2

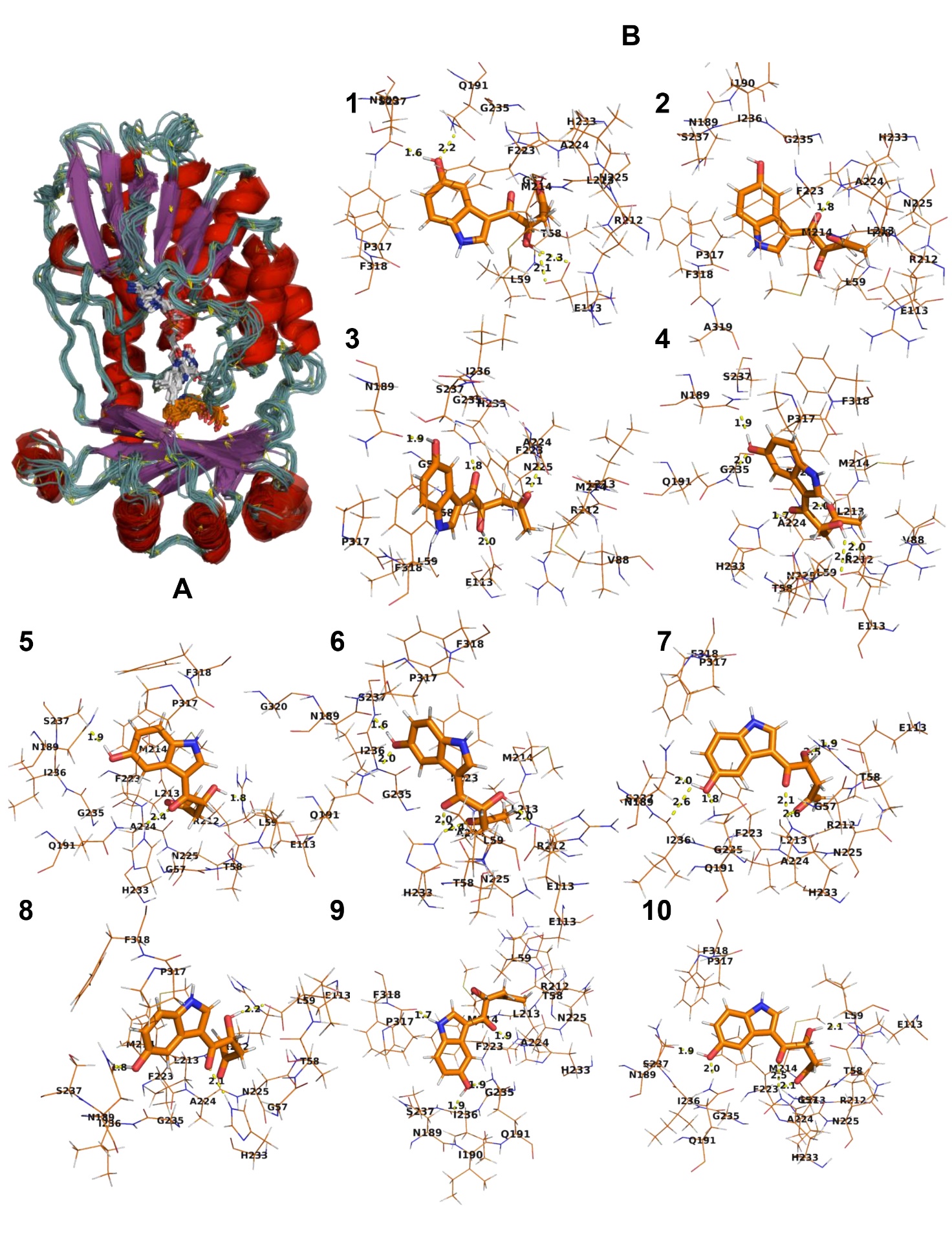

**Supplementary Table 1. Percentage of Structure coverage in the free energy landscape.**

| **Range** | **TDC (%)** | **CMNPD29406 (%)** | **CMNPD17967 (%)** | **CMNPD25139 (%)** |
| --- | --- | --- | --- | --- |
| Range 0-5  Range 5-10  Range 10-15  Range 15-20  Range 20-25  Range 25-30 | 84.71  11.40  2.44  0.92  0.43  0.06 | 89.84  8.23  1.66  0.25 | 84.09  11.26  3.03  1.24  0.35 | 87.89  10.6  1.43  0.05 |

**Supplementary Fig 5**. Aggregated motion of top ten principal components

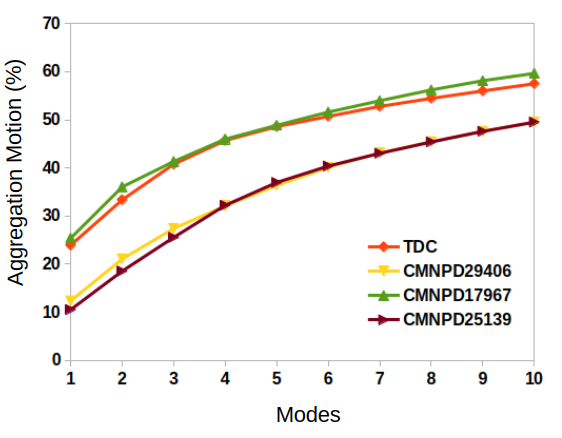
